# Virtual screening of MAP-*Tau* protein inhibitors from *Semecarpus anacardium* Linn. leaf extract for cancer prevention

**DOI:** 10.1101/2020.01.08.899708

**Authors:** Rajesh Kumar Singh, Anil Kumar Singh, Amit Ranjan, Akhileshwar Kumar Srivastava, Monika Singh, Abhishek Kumar, Kamal Nayan Dwivedi

## Abstract

*Semecarpus anacardium* is a well known Indian medicinal plant with various medicinal properties like hypoglycemic, antioxidant, anticancer, anti-inflammatory, anti-geriatric, antimicrobial and hair growth promoter, etc. The molecular mechanism of metabolites from fruiting bodies of *S. anacardium* against cancer has been described but anticancerous properties in its leaves are still unknown. The leaves were extracted in petroleum ether, ethyl acetate and methanol and assayed for anticancer activity using MTT assay. The active extract was evaluated for mode of cell death induction using EtBr-AO double staining and analyzed for phytochemical constituents using GC-MS, followed by molecular docking studies for exploration of possibility for anticancer agents and Drugability. In this study, ethyl acetate extract of leaf was found potent cytotoxic in MCF-7 cells and also induced apoptosis. It has also found the SLE is safe for normal cells. The molecular docking studies were done to explore the probable mechanism of action of the extract which showed 9 compounds are targeting the microtubule-associated protein tau (MAPT). MAPT promotes assembling and prevents disassembling to arrest the cell cycle. The overexpression of MAPT induces chemoresistance to cancerous cells against conventional drugs like paclitaxel. We have identified 17 compounds from ethyl acetate extract of *S. anacardium* leaves and drawn its chemical structure by using chembiodraw software to transform into pdb format. Further, the compounds have been subjected for molecular docking study to investigate its interactive efficiency with MAPT protein. The compound 13 had higher interactive potential to MAPT with binding energy −31.75 kcal/mol and lowest binding energy (−15.44 kcal/mol) was observed in compound 6. The present study suggested that the compounds from leaves of *S. anacardium* could be alternative approach of conventional drug for cancer treatment with cost effective and less side effect.

## INTRODUCTION

The microtubule-associated protein tau (MAPT) induces to assembling and disassembling of microtubule [1,2]. It is a tangle protein that interweaves α- and β-tubulin heterodimers in microtubule existing six isomeric forms [3,4] and binds on either side of microtubule surfaces exteriorly and interiorly to intact the structure of microtubule. The MAPT expresses in neurons [5] and lesser in other tissues [3] like liver [6], prostate [7], lungs [8], breast [9], pancreas [10], heart [11], skeletal muscle [12], kidney [13], etc.

Although, over expression of MAPT, alters the normal cellular function via enhancing microtubule polymerization and diminishing cell flexibility [14]. Microtubules are intracellular dynamic cytoskeletal polymers that play a vital role in several eukaryotic cellular processes including cell growth, division, mobility, shape and intracellular trafficking [15,16]. Dynamic microtubules are essential in mitotic cell division for proper attachment and segregation of chromosomes [17]. Microtubule is one of the significant target sites of Paclitaxel (PTX) [18]. It is widely used anticancer drugs in clinical practices in the treatment of breast, ovarian, cervical, endometrial, lung, gastric and other cancer [3].

It inhibits depolarization process of microtubules leading to conversation of unstable and dynamic microtubule into stable microtubule [15]. The stable microtubules are unable to spindle division, leading to cell cycle arrest in G2/M phase of mitosis and followed by apoptosis induction by another pathway, which collectively results into cell death [19–21]. As PTX binds on β-tubulin at inner surface of microtubules [3] and MAPT also binds on the same site at inner surface [22]. Owing to this, MAPT competes with PTX which leads the chemoresistance development among cancer cells against PTX [9,23]. Clinically, the patients with PTX chemoresistance show over expression of MAPT significantly [24]. Identification of MAPT inhibitors and their use in cancer treatment along with PTX improve the treatment results in the either case, PTX sensitive and resistant tumor [25,26].

*Semecarpus anacardium* Linn. (Family: Anacardiaceae) is a medicinal plant used in Ayurveda, a traditional medicine system of India for treatment of several diseases such as inflammation, neural problem, diabetese, cancer, geriatric problem, microbial infection and baldness [27]. It has been used as neurotonic formulation since a time ago in Ayurveda [28]. It have been reported by scientific study recently to posse various properties like hypoglycemic, antiatherogenic, antimicrobial, antioxidant, anticarcinogenic, antiinflammatory, anti-reproductive, CNS stimulant, skin diseases and hair growth promoter [28–33]. It shows in vitro antioxidant properties and induces cytotoxicity in liver, lung [34,35], breast, leukaemia, cervical cancer cells [36,37] through induction of apoptosis [37,38]. It also has antitumor activity in tumor bearing rat [39,40] and mice model [41]. The main chemical constituents are anacardic acid, bhilawanols, cardol, semecarpol and anacardol [28] in which bhilawanol and anacardic acids are responsible for the irritation, contact dermatitis, blisters and toxicity [42].

Virtaul screening and predicting drugs target using bioinformatics algorithm is an important tool for discovery and development of new drugs based on their molecular structure [43,44][45]. It is widely applied for hypothesize, evaluation of affinity with receptor-ligand interaction, optimization of ligand molecule, pharmacophore identification, development of computational drug model and molecular docking studies. It is also used for theoretical studies of absorption, distribution, metabolism and excretion as well as toxicity of drug or molecule of interest [46][47].

In this study, we have taken the compounds identified in ethyl acetate fraction of *Semecarpus anacardium* leaf extract as hypothesized drug candidates for computational studies to explore their exclusive structural characters. We also have executed computational study to identify the possible targets, evaluate inhibitory potential of the compounds against their targets. Here, we selected MAPT as target molecule along with nine MAPT targeting compounds for this computational study. Fruits of the plant have reported for their medicinal values, although, there is lack of information about the usage of the leaves and even no computation study were found using these compound previously. The purpose of this study was to identify the potential MAPT inhibitors from phytochemicals for development of future adjuvant of PTX for cancer treatment.

## METHODS

### Plant Material Collection and Extraction

The leaves of *S. anacardium* Linn. were collected in the month of January, 2018 from Vindhya forest region of Mirzapur district, Uttar Pradesh, India and authenticated by Prof. Anil Kumar Singh, Department of Dravyaguna, Faculty of Ayurveda, Institute of Medical Sciences, Banaras Hindu University, Varanasi, India. The leaves were washed thrice with tap water, dried under the shade and grinded it into powder using domestic electric grinder. The powder of the leaves was macerated into petroleum ether, ethyl acetate and methanol respectively thrice in each. The filtrates were concentrated using Buchi rotary evaporator and further dried using desiccators. The extracts of petroleum ether, ethyl acetate and methanol were coded as BLPE, BLEA and BLM respectively.

### Cell lines culture

Human breast cancer (MCF-7) cells, human liver cancer (HepG2) cells, human colorectal cancer (HCT-15), mice ascites carcinoma (EAC) cells and mice fibroblast (NCTC clone 929, L929) cells were procured from National Centre for Cell Sciences (NCCS), Pune, India and maintained in Minimum Essential Medium [MEM(E)] containing 10% fetal bovine serum (FBS), 100 unit/mL of penicillin G, 100 μg/mL of streptomycin sulfate and 250 ng/mL amphotericin B were added. The cell lines were maintained in a humidified environment with 5% CO2 gas and 37 °C temperature. At about 70-80% confluence and log phase, cells were harvested by trypsinization and used for all experimental studies.

### Cytotoxicity assay

The cell toxicity of extracts in different cancer cells and mouse fibroblast cells were evaluated by the MTT (3-(4,5-dimethylthiazol–2-yl)-2,5-diphenyl tetrazolium bromide) assay[48]. Approximately, 1□×□10° cells/well were seeded into 96 well plates culture plate and incubated for 24 h. then the cells were treated with the extracts and further incubated for 72 h. Then media was replaced with 50□L of MTT solution at a concentration of 5 mg/ml and again incubated for 4 h. after incubation at 37 °C, then supernatant was replaced with 100□l DMSO to solubilize the formed violet colored crystals. Finally, the absorbance was measured using microplate reader (Biorad, India) at a wavelength of 570□nm and percentage cell viability was calculated using the formula:

Percentage of cell viability = (OD of the treated cells /OD of the untreated cells as control) × 100.

### Analysis of Cellular morphology and cell death

The cells were treated as previously described and then the mode of cell death was detected by double staining with Acridine orange (AO) and Ethidium bromide (EtBr) and washed 10 mM PBS (pH 7.4) twice. The cells were visualized by fluorescence microscopy using blue filter (Dewinter, India) at 400× magnification[49].

### Phytochemical Screening using GC-MS

The phytochemical screening was was carried out by GC-MS system (JEOL GCMATE II GC-MS) at Sophisticated Analytical Instrumnent facility, Indian Institute of Technology Madras, India using previously described protocols[50]. The data were matched with the NIST library to identify the compounds.

### Ligand preparation and Potential protein target identification

The structures of identified compounds (table-1) had drawn using ChemBioDraw 14.0, converted into three dimensions in protein data bank file format. The compounds were subjected to identify the potential protein target through the Swiss Target Prediction server (http://www.swisstargetprediction.ch) using its SMILES (Simplified molecular-input line-entry system) format of each compound and *Homo sapiens* was considered as the source of target [51].

### ADME screenings

The molecular properties of compounds used in formulating “rule of Lipinski’s five rule” states that the molecule with less than 500 Dalton molecular weight, 5 or less number of hydrogen bond donor, 10 or less hydrogen bond acceptor and log P not greater than 5 (good membrane permeability) would be a good drug candidate [45,52]. ADME/Tox (Absorption, Distribution, Metabolism, Excretion and Toxicity) assessment plays a vital role in drug development, which helps in detecting drug likeliness of the compound. The ADME/Tox screening of compounds was performed at the SwissADME server (http://www.swissadme.ch) using the SMILES format of each compounds at default parameters [47].

### Molecular docking of compounds

The molecular docking of compounds is a computational approach for drug designing and development in which the binding mode of compounds with receptor is assessed [53]. The binding mode of compounds with MAPT was investigated to determine the conservative residues and active sites. Docking study of each compound was carried out using PachDock online server (https://bioinfo3d.cs.tau.ac.il/PatchDock) using clustering RMSD 4.0 and complex type default as settings. The results of each compound with solution number, docking score, area, atomic contact energy and six-dimensional transformation space was obtained. The results obtained through PatchDock shows the ranking of solution according to their geometric shape complementarily score on the basis of molecular shape representation and surface patch matching. The obtained results were further refined using FireDock online server (http://bioinfo3d.cs.tau.ac.il/FireDock) and top 10 solutions out of 1000 rescored solution were considered for the studies which represented the energy involvement in complex formation as global binding energy, attractive van der Waals, and hydrogen bond energies, etc. Each ligand-receptor complex of FireDock was ranked according to their minimum global binding energy. Chimera 1.12 was used to view the surface of the complex and Discovery Studio 2017 R2 Client was used to determine the mode of interaction of the complex formed [52].

## RESULTS AND DISCUSSION

In this study, the *S. anacardium* leaf was investigated for anticancer properties first time, which could be used as alternative for therapeutic purposes. Here, the leaf extracts were prepared using petroleum ether, ethyl acetate and methanol and coded as SLP, SLE and SLM respectively. Further, it was evaluated for anticancer activity in different cancer cells along with normal fibroblast, L929 cells. SLE extract was found most potent in MCF-7 cell line, was selected for fluorescence microscopy for detection of mode of cell death.

### The extracts induce cytotoxicity in cancer cells

Among the three extracts, The petroleum ether extract (SLP) of the leaves was most effective in human breast cancer cells (IC_50_ value 4.18 μg/ml), followed by human colorectal cancer cells (IC_50_ value 7.35 μg/ml), mouse ascetic carcinoma cells (IC_50_ value 8.88 μg/ml) and very few toxic for normal fibroblast cells (IC_50_ value 33.28 μg/ml) and human liver cancer cells (IC_50_ value 66.78 μg/ml). The ethyl acetate leaf extract (SLE) was most cytotoxic in human breast cancer cells (IC_50_ value 1.09 μg/ml), followed by mouse ascetic carcinoma cells (IC_50_ value 2.13 μg/ml), human colorectal cancer cells (5.43 μg/ml) and very least toxic for normal fibroblast cells (IC_50_ value 30.28 μg/ml) and human liver cancer cells (IC_50_ value 44.93 μg/ml). Similarly, the methanolic leaf extract (SLM) showed very potent toxic for human breast cancer cells (IC_50_ value 2.4 μg/ml), followed by mouse ascetic carcinoma cells (IC_50_ value 12.07 μg/ml), human colorectal cancer cells (IC_50_ value 13.03 μg/ml), normal fibroblast cells (IC_50_ value 18.02 μg/ml) and least for human liver cells (IC_50_ value 46.79 μg/ml). Overall, the ethyl acetate extract of leaves was potent cytotoxic in human breast cancer cells and fairly safe for normal mouse fibroblast cells (Fig. 1A, 1B &1C). The cytotoxicity of leaf extracts is almost similar to the previous studies of nut extract of the same plant[36,54]. Moreover, SLE is safer than nut extract[37], therefore, based on the ratios IC_50_ values of extract in normal cells to cancer cells (selectivity Index), SLE has found the potent extract and selected for further studies.

**Figure 1:**
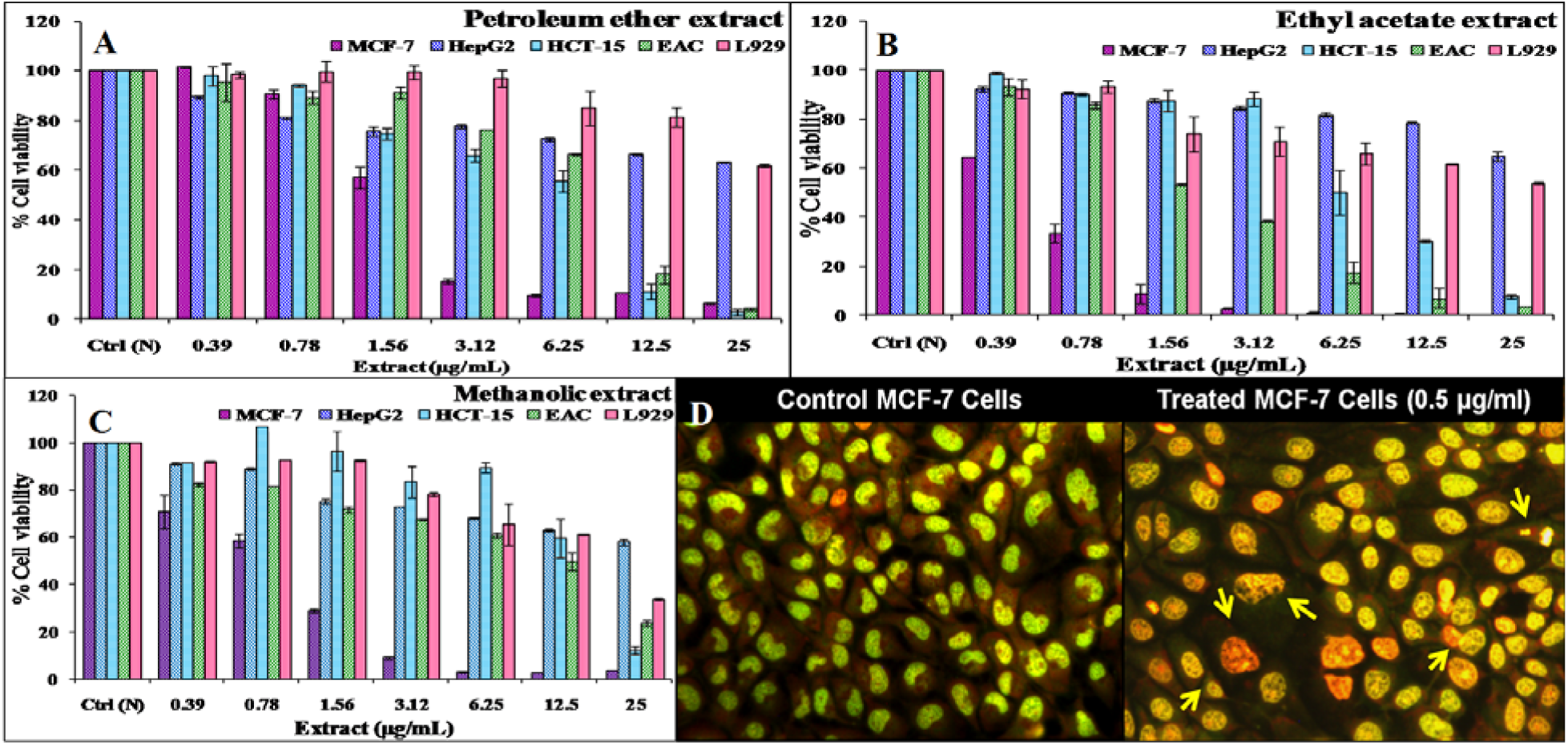
Cytotoxic effect of different extracts of *S. anacardium* leaf extract with their IC_50_ value (**A**) Petroleum ether extract (SLP): MCF-7 cells (4.18 μg/ml), HepG2 (66.78 μg/ml), HCT-15 cells (7.35 μg/ml), EAC cells (8.88 μg/ml) and L929 cells (33.28 μg/ml). (**B**) Ethyl acetat leaf extract (SLE): MCF-7 cells (1.09 μg/ml), HepG2 (44.93 μg/ml), HCT-15 cells (5.43 μg/ml), EAC cells (2.13 μg/ml) and L929 cells (30.28 μg/ml), and (**C**) Methanolic leaf extract (SLM): MCF-7 cells (2.4 μg/ml), HepG2 cells (46.79 μg/ml), HCT-15 cells (13.03 μg/ml), EAC cells (12.07 μg/ml) and L929 cells (18.02 μg/ml). (**D**) Fluorescence microscopy to evaluate cell death using Ethidium bromide-Acridine orange double staining in MCF-7 cells treated with SLE.

### SLE induces apoptosis in MCF-7 cells

The ethidium bromide is impermeant to the cells membrane, therefore binds with DNA stands only when the plasma membrane broken, leading to red color fluorescence, while Acridine orange is a cell membrane permeant dye, has affinity to bind with nucleic acids, leading to green fluorescence[55]. In combination of these two fluorescing dyes, yellowish orange color developed for early apoptosis, orange for apoptosis and red for late apoptosis or necrosis mode of cell death[56]. The fluorescence microscopic analysis after AO-EtBr double staining showed that SLE induced apoptosis in human breast cancer (MCF-7) cells as seen the orange-red fluorescence with nuclear fragmentation (Fig. 1D). The untreated cells showed intact nuclear morphology [57].

### GC–MS analysis of SLE and protein target identification

The GC-MS data analysis identified 17 compounds in SLE extract (table 1 and fig. 2). Intentionally, human MAPT protein (PDB ID: 2MZ7) was selected as target molecule for the identified seventeen compounds, in which nine compounds **(2, 5, 8, 9, 10, 14, 15, 16** and **17)** had potential protein target for the microtubule associated protein tau (MAPT). Now, it has been selected the nine MAPT targeting compounds for study on different parameters of drug-ability against human origin MAPT protein.

**Table 1:**
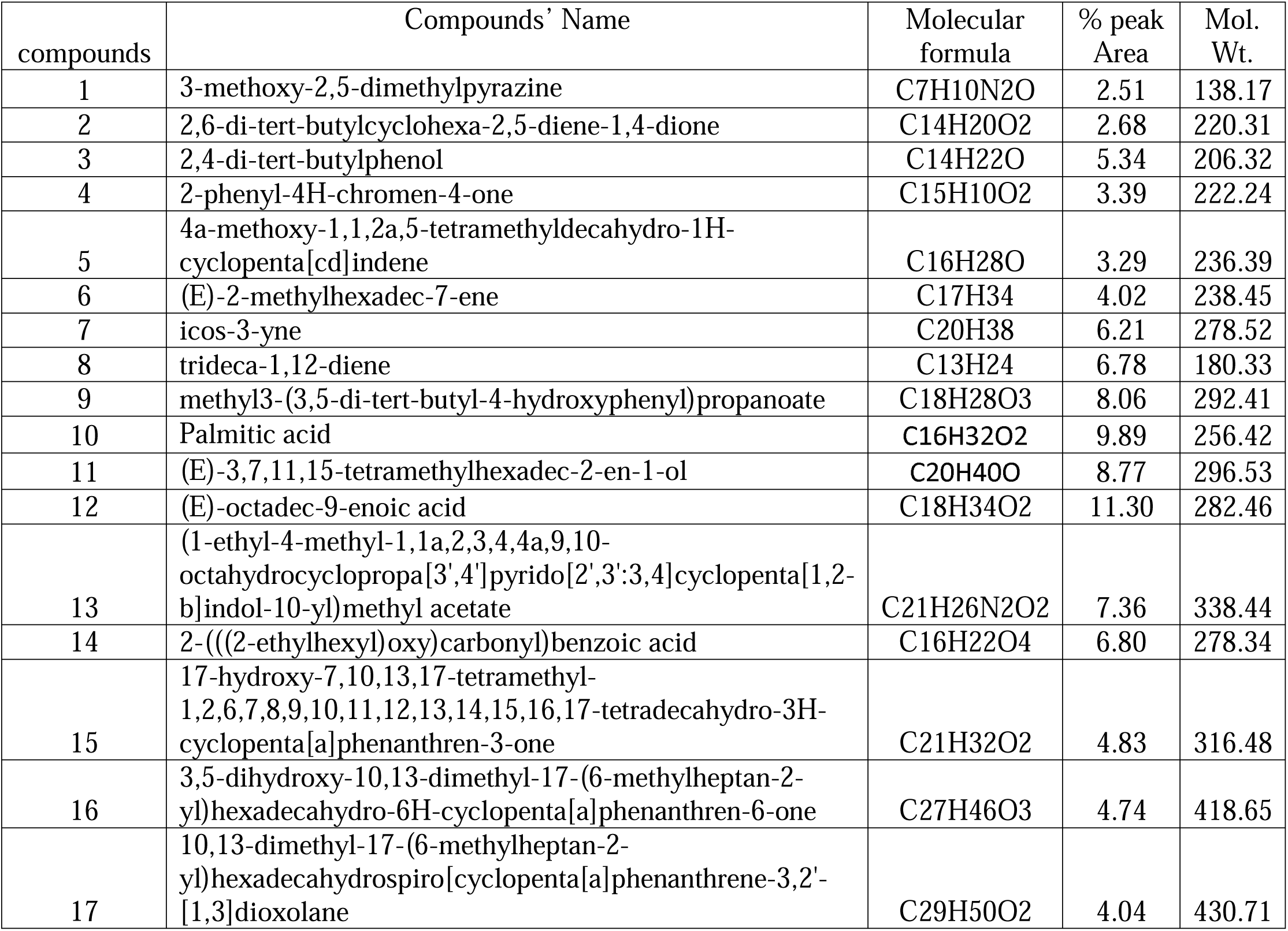
Chemical composition of ethyl acetate extract of *S. anacardium* leaves.

| compounds | Compounds' Name | Molecular formula | % peak Area | Mol. Wt. |
| --- | --- | --- | --- | --- |
| 1 | 3-methoxy-2,5-dimethylpyrazine | C7H10N2O | 2.51 | 138.17 |
| 2 | 2,6-di-tert-butylcyclohexa-2,5-diene-1,4-dione | C14H20O2 | 2.68 | 220.31 |
| 3 | 2,4-di-tert-butylphenol | C14H22O | 5.34 | 206.32 |
| 4 | 2-phenyl-4H-chromen-4-one | C15H10O2 | 3.39 | 222.24 |
| 5 | 4a-methoxy-1,1,2a,5-tetramethyldecahydro-1H-cyclopenta[cd]indene | C16H28O | 3.29 | 236.39 |
| 6 | (E)-2-methylhexadec-7-ene | C17H34 | 4.02 | 238.45 |
| 7 | icos-3-yne | C20H38 | 6.21 | 278.52 |
| 8 | trideca-1,12-diene | C13H24 | 6.78 | 180.33 |
| 9 | methyl3-(3,5-di-tert-butyl-4-hydroxyphenyl)propanoate | C18H28O3 | 8.06 | 292.41 |
| 10 | Palmitic acid | C16H32O2 | 9.89 | 256.42 |
| 11 | (E)-3,7,11,15-tetramethylhexadec-2-en-1-ol | C20H40O | 8.77 | 296.53 |
| 12 | (E)-octadec-9-enoic acid | C18H34O2 | 11.30 | 282.46 |
| 13 | (1-ethyl-4-methyl-1,1a,2,3,4,4a,9,10-octahydrocyclopropa[3',4']pyrido[2',3':3,4]cyclopenta[1,2-b]indol-10-yl)methyl acetate | C21H26N2O2 | 7.36 | 338.44 |
| 14 | 2-(((2-ethylhexyl)oxy)carbonyl)benzoic acid | C16H22O4 | 6.80 | 278.34 |
| 15 | 17-hydroxy-7,10,13,17-tetramethyl-1,2,6,7,8,9,10,11,12,13,14,15,16,17-tetradecahydro-3H-cyclopenta[a]phenanthren-3-one | C21H32O2 | 4.83 | 316.48 |
| 16 | 3,5-dihydroxy-10,13-dimethyl-17-(6-methylheptan-2-yl)hexadecahydro-6H-cyclopenta[a]phenanthren-6-one | C27H46O3 | 4.74 | 418.65 |
| 17 | 10,13-dimethyl-17-(6-methylheptan-2-yl)hexadecahydrospiro[cyclopenta[a]phenanthrene-3,2'-[1,3]dioxolane | C29H50O2 | 4.04 | 430.71 |

**Figure 2:**
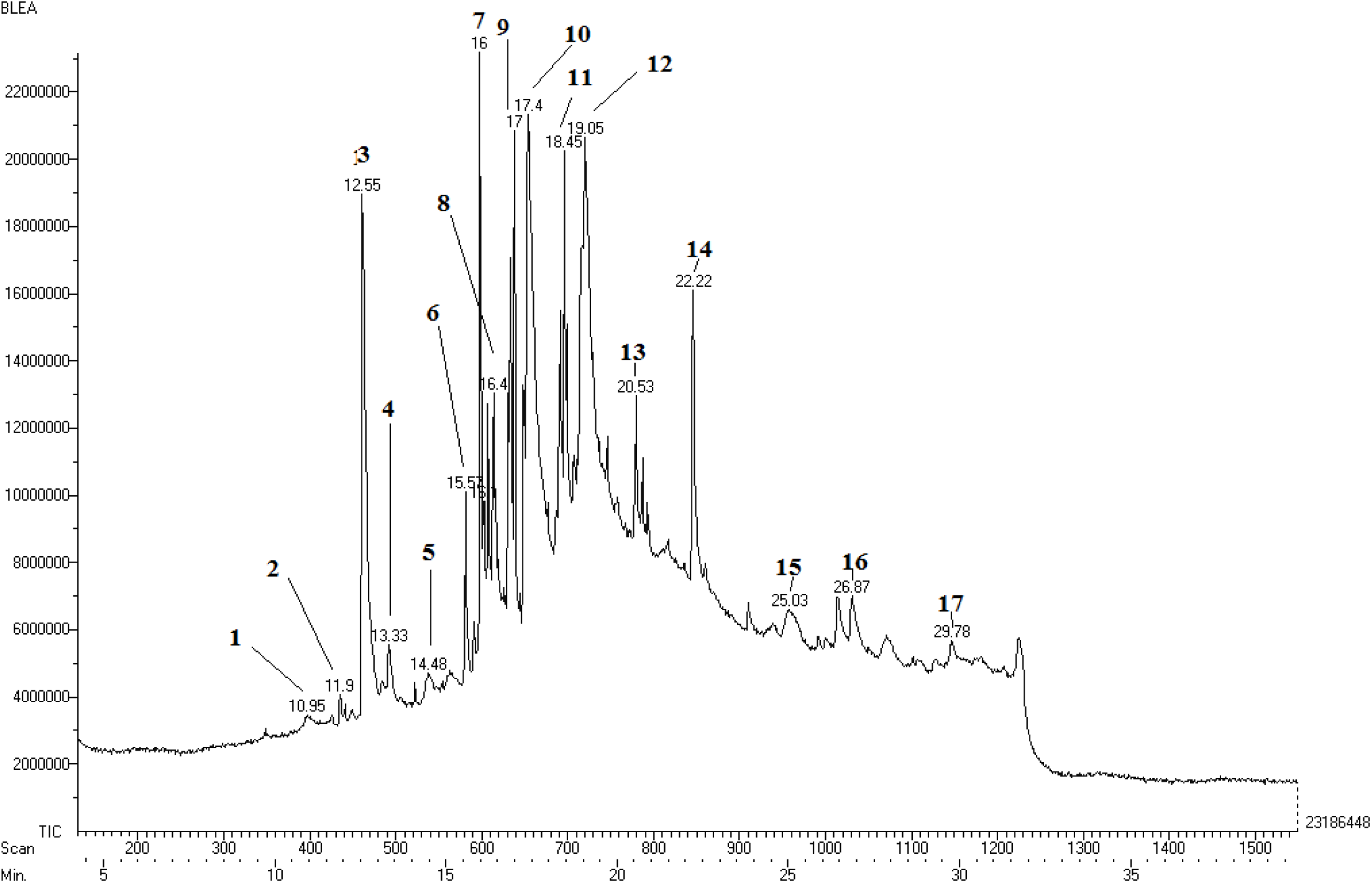
GC-MS spectral chromatogram *Semecarpus anacardium* leaf extract (SLE).

### Molecular Docking studies

The interactive residues of target protein and ligand help the researchers to design and develop the drugs that are more efficient and specific for their target protein. The docking of MAPT target protein with compounds revealed the involvement of H-bonding between ligand and amino acids residues of MAPT along with other non-covalent interactions such as hydrophobic interactions and van der Waals forces which play an important role in formation of complexes. The surface view of complex reveals that the binding pocket of tau-protein with area 287.30 Å was made for accommodation of **compound 2** (Fig.3.1A). Fig. 3.1B and 3.1C represented the 3D/2D of complexed structure showing the binding mode of **compound 2** with residues (ILE297, ASN296, LYS290, HIS299, CYS291, GLY292, and SER293) of tau-protein indicating that the **compound 2** is capable to disrupt the natural integrity of tau-protein. **Compound 5** fitted on the binding pocket (251.90 Å) of tau-protein (Fig. 3.2A) and interacted with residues (LYS267, LYS280, LYS281 and LEU284) of tau-protein showing in 3D/2D structure (Fig. 3.2B and 3.2C). It has suggested that the **compound 5** may disturb the oncogenic activity of tau-protein. The complexation of **compound 8** on the surface of tau-protein with area 285.90 Å (Fig. 3.3A) was made through interaction of residues (VAL287, LYS294, ILE308, TYR310 and PRO312) of MAPT (Fig. 3.3B and 3.3C) may alter the function of tau-protein. The binding pocket area (335.20 Å) of the complex is made for accommodation of **compound 9** (Fig. 3.9A). The complex shows interaction with residues (SER293, LYS294, ILE297, LYS298 and ILE308) of MAPT (Fig 3.9B and 3.9C) that may down regulate the activity of tau protein in cancer. The binding pocket (344.60 Å) of tau-protein with area 344.60 Å (Fig. 3.4A) made complex by accommodating of **compound 10** through different residues (ILE278 and VAL287) of tau-protein (Fig. 3.4B and 3.4C) and the made complex may inhibit the function of tau-protein MAPT function. The binding pocket of tau-protein with area 312.00 Å integrated with **compound 14** as visualized by surface view of complex (Fig. 3.5A). Fig. 3.5B and 3.5C represented complex with binding mode of **compound 14** with different residues (HIS268, SER285, GLN288 and SER289) of tau-protein and this complex may interrupt the structure of tau-protein. The **compound 15** was also attached with MAPT surface with pocket area 285.40 Å (Fig. 3.6A) by interacting with residue HIS268 of tau-protein that may change natural confirmation of tau-protein (Fig. 3.6B). The surface view of binding pocket (418.90 Å) of MAPT complexed with **compound 16** (Fig. 3.7A) by integrating with residues (LEU284, CYS291 and ILE297) of MAPT (Fig. 3.7B and 3.7C. The interacted **compound 16** might disrupted the integrity of MAPT. Figure 3.8A exhibits the binding pockets of MAPT for **compound 17** with pocket area 453.10 Å. The compound revealed interactions with two amino acid residues (ILE277 and CYS291) ((Fig. 3.8B and 3.8C) indicating that the **compound 17** has capability to suppress of oncogenic activity of MAPT. The paclitaxel accommodated into binding pocket of MAPT with area 343.20 Å (Fig. 3-PTX-A). Fig. 3-PTX-B and 3-PTX-C represented the 3D and 2D structure of the complex showing the binding mode of paclitaxel with residues (HIS268, GLN269, GLY271, ASP283 and ASN286) of tau-protein that may also alter integrity of tau-protein.

**Figure 3:**
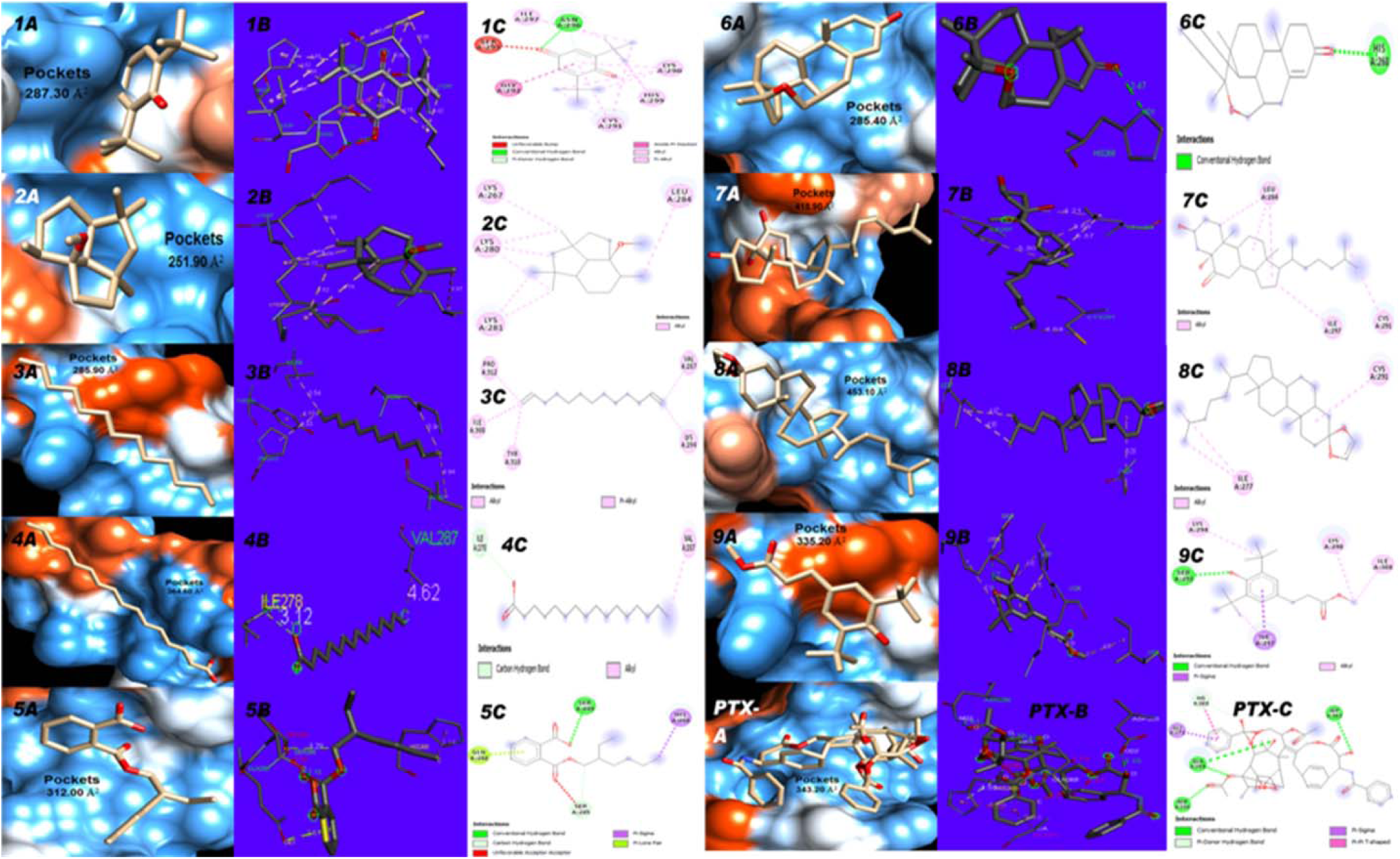
Visualization of the interaction of receptor protein, MAPT with ligands molecules in which A represents 3D surface view of binding site of compounds, B represents3D spacial interactions and C represents 2D interaction of the compounds respectively. 1, 2, 3, 4, 5, 6, 7, 8 and 9 represents **compound 2, 5, 8, 9, 10, 14, 15, 16** and **17** respectively.

**Figure 4:**
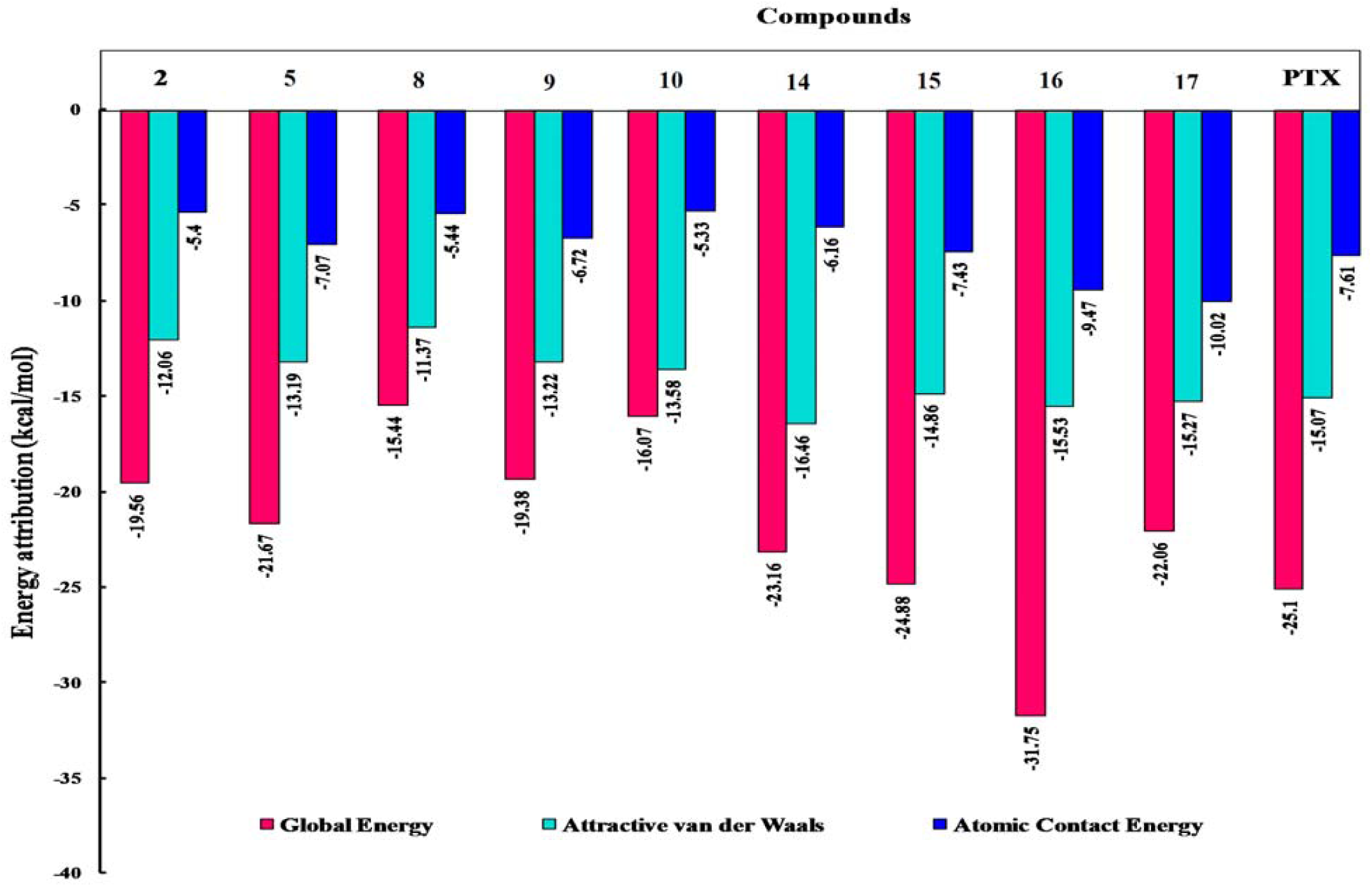
Representing the inherent free binding energy (kcal/mol) attributes of the complex of the compounds with MAPT.

### Analysis of binding energy in complexes

The interactions (van der Waals interaction, hydrogen bond, alkyl bond, etc) of receptor-ligand and their potential energy determine the fate of the ligand molecule. These interactions contributed to stable bond formation between ligands and protein structure (Fig. 4). The global energy of the protein-ligand complex is directly associated with stability of the complex and its lower value indicates higher interaction probability [58][45]. The molecular docking analysis of compounds against target protein, MAPT revealed that the compound 16 (−31.75 kcal/mol) able to establish more stable bonds than conventional drug paclitaxel (−25.10 kcal/mol) whereas compound 15 (−24.88 kcal/mol), compound 14 (−23.16 kcal/mol), compound 17 (−22.06 kcal/mol), compound 5 (−21.67 kcal/mol), compound 2 (−19.56 kcal/mol), compound 9 (−19.38 kcal/mol), compound 10 (−16.07 kcal/mol) and compound 8 (−15.44 kcal/mol) had higher global energy, i.e. less stable bonds respectively. The van der Waals interaction plays an important role in complex formation [59] which indicates that the more stable bond with the more negative value of attractive van der Waals and Atomic contact energy. These interactions are collectively contributes in global energy [60,61].

## CONCLUSION

In this study, we have identified 17 compounds in the ethyl acetate extract of *S. anacardium* leaves in which **compounds 2, 5, 8, 10, 14, 15, 16, 17** and **17** have good MAPT binding potency. The compound **compounds 16** has higher interaction potential to MAPT (−31.75 kcal/mol) and **compounds 8** has lowest binding potency (−15.44 kcal/mol). Compound **8, 9, 10** and **14** are major compound in the extract whereas compound **14** and **15** compete with paclitaxel to accommodate at amino acid HIS268 of MAPT. This study suggests that the compounds of *S. anacardium* could be alternative approach of conventional drug for cancer treatment with cost effective and less side effect. Though, the limitations of this computational study were associated with the reliability in *in vivo* study. As *in silico* studies might be less reliable in *in vivo* studies. However, our computational studies suggested that the identified compounds may play an important pharmacological role in chemotherapy for paclitaxel chemoresistance cancer cells through targeting MAPT. Hence, the compounds could be alternative to conventional drugs in prevention of tumor progression in cancer patients.

## Author Contributions

RKS, RT, AKS and SKS designed the study; RKS performed the experimental and docking studies; RKS, AR, MS, KND, AK, AKS and SKS contributed to the formatting, writing and revision of the manuscript. All authors have read and approved the manuscript for publication.

## Acknowledgement

We are thankful to the Sophisticated Analytical Instruments Facility, Indian Institute of Technology Madras, Chennai and Experimental facilities, Centre of Experimental Medicine & Surgery (CEMS), Institute of Medical Sciences, Banaras Hindu University, Varanasi, India for instrument facilities. RKS, AR and RT are grateful to BHU and UGC for financial supports as fellowships.

## Abbreviations

MAPT: Microtubule associated protein-*Tau*.
GC-MS: Gas chromatography-Mass spectroscopy
G2/M: Gap 2/Mitosis phase.
PTX: Paclitaxel.
PDB: Protein data bank.

